# Extracellular vesicles produced during fungal infection in humans are immunologically active

**DOI:** 10.1101/2024.03.20.585987

**Authors:** Caroline P. de Rezende, Patrick W. S. Santos, Renan A. Piraine, Virgínia C. Silvestrini, Julio C. J. Barbosa, Fabiana C. P. Valera, Edwin Tamashiro, Guilherme G. Podolski-Gondim, Silvana M. Quintana, Rodrigo Calado, Roberto Martinez, Taicia P. Fill, Márcio L. Rodrigues, Fausto Almeida

## Abstract

Of the known 1.5 million fungal species, *Candida* spp., *Cryptococcus* spp., and *Paracoccidioides* spp. are the main pathogenic species causing serious diseases with almost two million annual deaths. The diagnosis and treatment of fungal infections are challenging since of the limited access to diagnostic tests and the emergence of antifungal resistance. Extracellular vesicles (EVs) promote the interactions of fungal cells with other organisms and play an important role in the pathogen–host relationship. Owing to the complexity of fungal EVs and the lack of clinical studies on their roles in human infections, we studied the EVs from the serum and urine samples of patients with fungal infections caused by *Candida albicans*, *Cryptococcus neoformans*, and *Paracoccidioides brasiliensis* and determined their roles. Steroids, sphingolipids, and fatty acids were identified as the main secondary metabolites via mass spectrometry analysis. We asked whether these metabolites in EVs could play roles in modulating the host immune response. Our findings revealed the polarization of the proinflammatory profile in murine and human macrophages, with the increased production of cytokines, such as the tumor necrosis factor-α, interferon-γ, and interleukin-6, and an increased expression of the inducible nitric oxide synthase gene, a M1 response marker. Therefore, circulating EVs from patients with fungal infections are likely involved in the disease pathophysiology. Our findings provide insights into the roles of EVs in fungal infections in clinical samples and in vitro, suggesting possible targets for systemic mycoses therapy.

**Significance Statement:** Fungal infections cause approximately 1.6 million deaths annually. Due to therapeutic and diagnostic limitations, it is mandatory to understand and develop new immunological interventions. Despite several in vitro studies on the production of extracellular vesicles (EVs) from fungal pathogens, this study is a pioneer in the identification and characterization of EVs in the course of fungal infection in humans. Our group demonstrated the presence of EVs in clinical samples from patients diagnosed with candidiasis, cryptococcosis, and paracoccidioidomycosis, as well as the EVs interaction produced by host and fungal pathogen with the immune system, resulting in relationships that may be beneficial for the progression or elimination of fungal disease.

## Introduction

High incidence of fungal infections and their therapeutic limitations necessitate the in-depth study of pathogenic fungi to develop new immunological interventions (1). The World Health Organization classifies fungal pathogens into three priority groups: critical, high, and medium. *Candida albicans* and *Cryptococcus neoformans* are critical priority fungi, whereas *Paracoccidioides brasiliensis* is a medium priority fungus (2). Currently, most common fungal infections are invasive candidiasis (70%), cryptococcosis (20%), and aspergillosis (10%) (3). The emergence of fungal species resistant to the four classes of antifungal drugs (azoles, polyenes, echinocandins, and flucytosine) poses a serious challenge for the effective treatment of fungal infections (4, 5).

High mortality rates of pathogenic fungal infections are due to underdiagnosis and limited treatment options. Emergence of multidrug-resistant fungal pathogens necessitates the rapid and accurate diagnosis of causative fungi for effective treatment (6). Diagnostic tests, including the gold standard test, have various limitations, such as low sensitivity, invasive collection process, need for qualified professionals, and slow response time, emphasizing the need for new microbiological diagnostic tools with greater specificity and sensitivity for the detection and treatment of patients in clinical settings (7, 8).

Biological fluids, such as blood and urine, are important sources of extracellular vesicles (EVs), providing a promising platform for the identification of diagnostic biomarkers (9, 10). EVs consist of spherical lipid bilayers, whose composition depends on the origin of the cell and environmental and metabolic conditions. They carry lipids, carbohydrates, proteins, and nucleic acids (11). Moreover, EVs facilitate bidirectional communication between the pathogen and host, influencing the host immune responses (12). Monocytes and dendritic cells are modulated by fungal EVs (13, 14), such as those derived from the fungal pathogens, *C. albicans*, *C. neoformans*, and *P. brasiliensis*, which change the macrophage functions in vitro by modulating the host immune response during infection (15, 16).

Despite many studies on the production of EVs by fungal pathogens (15, 17–19), to date, no studies have characterized the roles of EVs in fungal infections using human samples. Here, we identified the presence of EVs in the serum and urine samples of patients with candidiasis, cryptococcosis, and paracoccidioidomycosis, suggesting the potential of fungal EV–host interactions for the development of new therapeutic strategies. Identification of secondary metabolites via mass spectrometry (MS) analysis revealed an abundance of lipids, especially steroids, sphingolipids, and fatty acids, suggesting their crucial roles in the physiological functions of EVs. Furthermore, EVs derived from infected patients polarized the human (THP-1) and murine (AMJ2-C11) macrophages towards a proinflammatory immune response profile in vitro. Taken together, our results suggest that both the host- and pathogen-derived EVs are involved in the eradication of fungal infections via modulation of the host immune response. Therefore, our study provides new insights into the modulatory mechanisms of EVs and their potential use in novel therapeutic approaches for systemic mycoses.

## Results

### Identification and characterization of EVs from the serum and urine samples of patients with fungal infections

In this study, serum and urine samples of patients infected by *C. albicans* (n=10), *C. neoformans* (n=6), *P. brasiliensis* (n=20), and healthy individuals (n=20) were collected for the identification and characterization of EVs via nanoparticle tracking analysis (NTA; NanoSight; NS300). The purified samples were randomly grouped into pools of five samples each from patients with candidiasis (Figure 1A and 1B) and paracoccidioidomycosis (Figure 1E and 1F), and a pool of six samples from patients with cryptococcosis (Figure 1C and 1D). Healthy individuals (n=20) were randomly grouped into a pool of five samples and used as non-infected controls (Supplementary 1). EV concentrations in the pools of serum samples from infected patients were 2.94×10^11^ EVs/mL (mean: 255 nm; mode: 196 nm) for candidiasis (Figure 1A), 3.29×10^12^ EVs/mL (mean: 150 nm; mode: 108 nm) for cryptococcosis (Figure 1C), and 1.7×10^12^ EVs/mL (mean: 215 nm; mode: 197 nm) for paracoccidioidomycosis (Figure 1E). Additionally, urine samples showed mean vesicle concentrations of 4.14×10^10^ EVs/mL (mean: 211 nm; mode: 112 nm) for candidiasis (Figure 1B), 3.26×10^10^ EVs/mL (mean: 131 nm; mode: 87 nm) for cryptococcosis (Figure 1D), and 6.95×10^10^ EVs/mL (mean: 236 nm; mode: 123 nm) for paracoccidioidomycosis (Figure 1F). Therefore, patients with candidiasis exhibited lower concentrations of larger sized EVs in the serum than those with cryptococcosis and paracoccidioidomycosis. These results demonstrate the presence of EVs in the serum and urine samples of patients with fungal infections, mainly those caused by *C. albicans*, *C. neoformans*, and *P. brasiliensis*.

**Figure 1.**
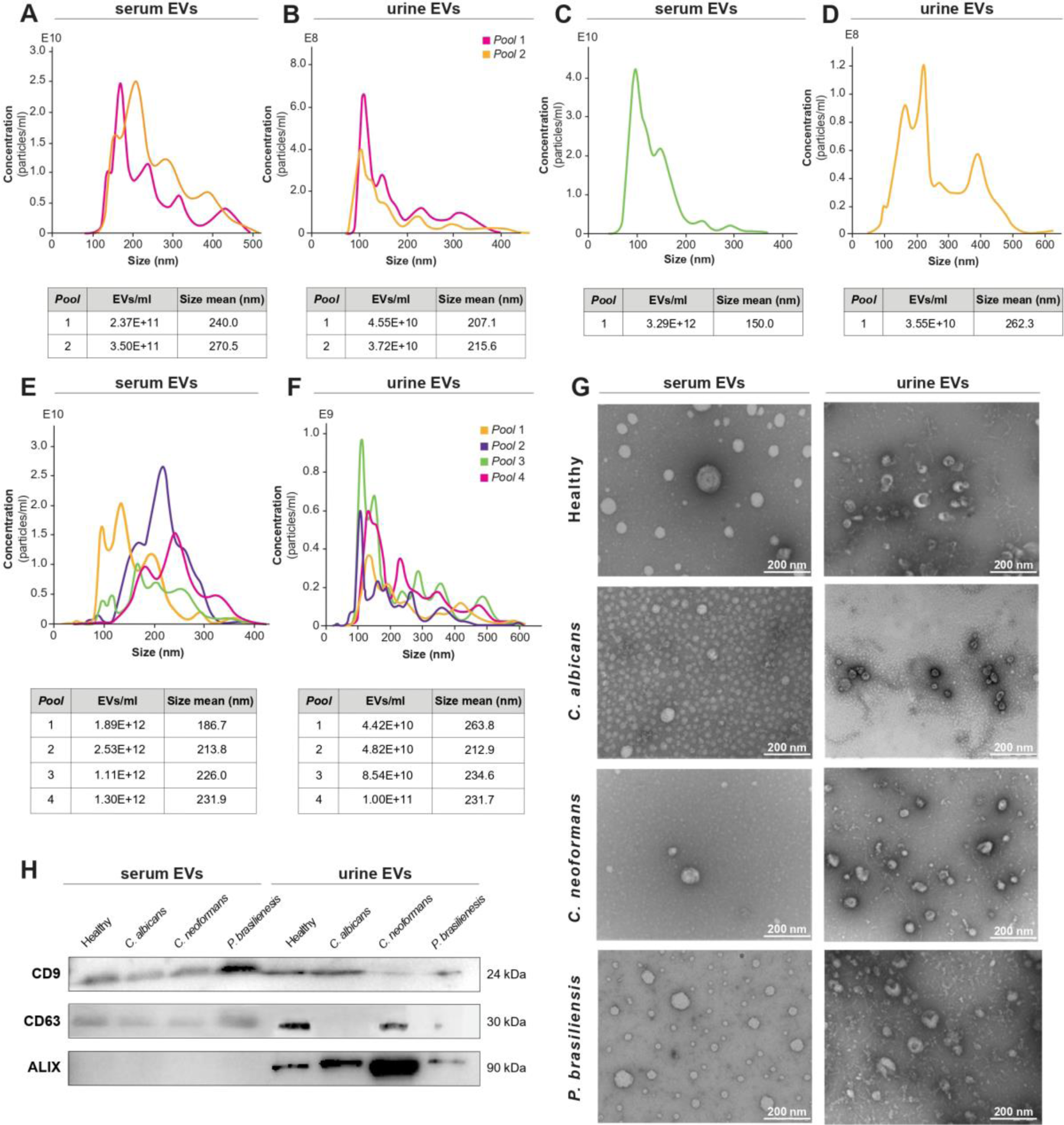
Characterization of extracellular vesicles (EVs) from the clinical samples of patients with fungal infections caused by *Candida albicans*, *Cryptococcus neoformans*, and *Paracoccidioides brasiliensis*. Nanoparticle tracking analysis (NTA) of the concentration (EVs/mL) and size (nm) of EVs from the serum and urine samples of patients with candidiasis (A and B), cryptococcosis (C and D), and paracoccidioidomycosis (E and F) from the Hospital das Clinicas, Ribeirao Preto Medical School (HC-FMRP). (G) Transmission electron microscopy was conducted on the clinical samples of healthy individuals and patients with fungal infections, and EVs with classic cup-shaped morphology were detected. Scale bar: 200 nm. (H) Western blotting of the protein extract from EVs derived from infected and non-infected patients with antibodies specific to the tetraspanin family (CD9 and CD63) and ALIX. Glyceraldehyde 3-phosphate dehydrogenase (GAPDH) was used as a loading control.

Spherical shape of EVs allows the calculation of their diameter using various techniques, such as transmission electron microscopy (TEM) that shows the resolution in nanometers (20). The classic cup-shaped morphology of EVs can be used to distinguish different classes of vesicles in the same samples (21, 22). The presence of EVs in the serum and urine samples of healthy individuals and patients with candidiasis, cryptococcosis, and paracoccidioidomycosis was visualized using TEM. The capture fields from TEM images revealed that EVs exhibited the classic cup-shaped morphology with a diameter of 200 nm (Figure 1G), thus confirming the characteristics of EVs from the serum and urine samples of patients with fungal infections.

### EVs from the serum and urine samples of patients with candidiasis, cryptococcosis, and paracoccidioidomycosis present classic EV markers

For the characterization of EV proteins, the International Society of Extracellular Vesicles (ISEV) recommends the evaluation of the expression of at least one protein from the following classes: transmembrane, anchored in glycosylphosphatidylinositol associated with plasma membranes and/or cytosolic, and heat shock proteins (23). Here, we analyzed the presence of three classic protein markers, CD9, CD63, and ALIX, in the serum and urinary EVs of healthy individuals and patients with fungal infections caused by *C. albicans*, *C. neoformans*, and *P. brasiliensis* using western blotting (Figure 1H). For CD9 and CD63 labeling, tetraspanin family, which consists of membrane proteins regulating the selective trafficking of associated proteins possibly via membrane compartmentalization, is crucial for EV characterization (24, 25). Here, we performed ALIX labeling of EVs from the urine samples of healthy individuals and patients with fungal infections (Figure 1H). Notably, EVs isolated from the clinical samples were characterized according to the standards proposed by ISEV.

### Statistical analysis and molecular networking in the serum and urinary EVs of healthy individuals and patients with fungal infections

EVs from the serum and urine samples of healthy individuals and patients with candidiasis, cryptococcosis, or paracoccidioidomycosis were prepared as dry extracts of lyophilized EVs and resuspended in MeOH at a concentration of 1 mg/mL for UHPLC-MS/MS analysis. The data obtained were analyzed using principal components analysis (PCA). The main components of PCA showed a clear separation between the analyzed serum and urinary EVs, corresponding to 37.9% of the total data variance, 23.4% for PC1, and 14.5% for PC2 (Figure 2A). This distinctive separation was fundamental to distinguish metabolic patterns associated with different health state conditions. Feature Based Molecular Networking (FBMN) was applied to analyze the chemical composition of the EVs under the studied conditions (Supplementary 2). The fragmentation tolerance parameters of the library data were 0.02 Da and the cosine score for the chromatograms that presented corresponding peaks was 0.7 (70%) (Supplementary 3). The annotation resulted in a total of 80 compounds (Annotation levels I and II) using the MS/MS libraries from GNPS and analytical standards. Among these, notable metabolites from the classes of steroids, sphingolipids, and fatty acids were annotated (Figure 2B).

**Figure 2.**
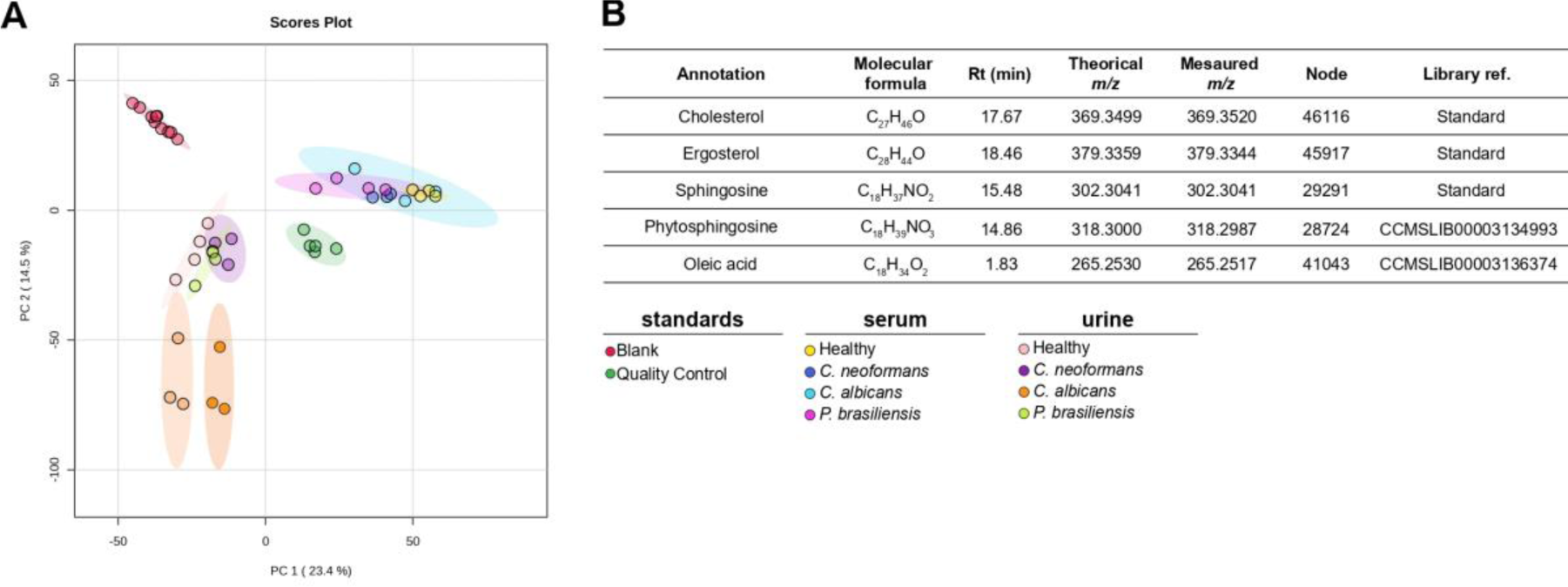
Statistical analysis of the metabolomics data of EVs from the serum and urine samples of healthy individuals and patients with fungal infections. (A) Data analysis using principal components analysis (PCA). (B) Annotation of the chemical compounds identified in EVs under experimental conditions using tandem mass spectrometry (MS/MS) libraries, with representation of the molecular formula, retention time (Rt), theoretical and measured *m/z*, node, and reference library.

### Metabolite annotations of serum EVs in patient samples

Metabolomics is a systems biology approach that aims to profile metabolites in cells, biofluids, and tissues (26). In the present study, the profiles of secondary metabolites in serum EVs from patients infected with *C. albicans*, *C. neoformans*, and *P. brasiliensis* were analyzed using UHPLC-HRMS/MS analysis, and serum EVs from healthy individuals were used as non-infection controls (Figure 3). The detection of cholesterol and ergosterol, which are compounds found in cell membranes, is important for understanding the origin of EVs and the relationship between pathogens and hosts. Cholesterol (C_27_H_46_O) was predominantly found in serum EVs from healthy individuals (Figure 3A), with a peak area of 1.1×10^6^, and with lower abundance in EVs from patients with fungal infections (Figure 3B), measured at *m/z* 369.3520 [M+H-H_2_O^]+^, and high similarity with the spectral library (Supplementary 3A). Ergosterol (C_28_H_44_O), measured at *m/z* 379.3344 [M+H-H_2_O]^+^ was detected in the serum EVs of patients with fungal infections and was not present in healthy individuals. EVs of *C. albicans* (peak area of 2.4×10^3^) and *C. neoformans* (peak area of 1.45×10^3^) showed a significant difference compared to EVs derived from healthy individuals (Figure 3, C and D; Supplementary 4).

**Figure 3.**
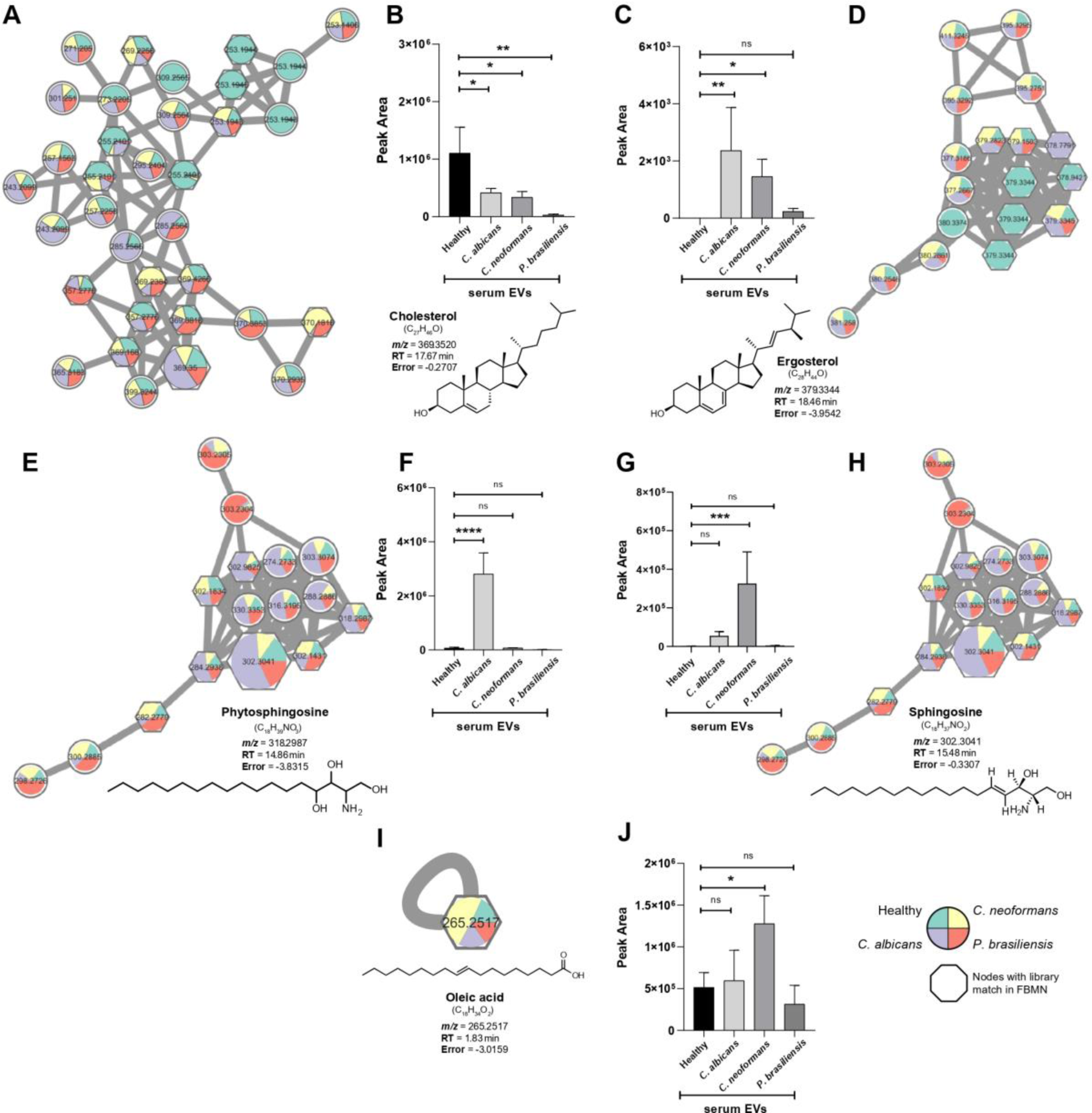
Annotations of secondary metabolites in the serum EVs of healthy individuals and patients infected by *C. albicans*, *C. neoformans*, and *P. brasiliensis*. Molecular networks generated by the GNPS platform (nodes with library correspondence in FBMN) and representation of the chemical structure of compounds belonging to the steroids (cholesterol and ergosterol) (A, D), sphingolipids (phytosphingosine and sphingosine) (E, H), and fatty acids (oleic acid) (I) classes. Statistical determination of the peak areas of steroids (B, C), sphingolipids (F, G), and fatty acids (J) in the serum EVs of healthy individuals and patients infected by *C. albicans*, *C. neoformans*, and *P. brasiliensis*. Healthy individuals were used as controls. Statistical difference: ns, p > 0.05; *p < 0.05; **p < 0.01; ***p < 0.001; ****p < 0.0001.

In EVs from the serum of patients with candidiasis and cryptococcosis, we identified compounds such as phytosphingosine (C_18_H_39_NO_3_, *m/z* 318.2987 [M+H]^+^) and identified sphingosine (C_18_H_37_NO_2_, *m/z* 302.3041 [M+H]^+^; Figure 3, E–H) using an analytical standard. These compounds are present in the sphingolipid pathway, with phytosphingosine being an intermediate metabolite and sphingosine being an 18-carbon amino alcohol derived from the cleavage of ceramide process (27). The peak area corresponded to 2.8×10^6^ for phytosphingosine (Figure 3F) and 5.4×10^4^ for sphingosine (Figure 3G) in serum EVs from patients infected by *C. albicans*. Furthermore, in serum EVs from patients infected with *C. neoformans*, the peak area was 3.25×10^5^ for sphingosine.

Oleic acid (cis-9-octadecenoic acid), commonly known as omega-9 fatty acid, has 18 carbons with a cis double bond at the 9th carbon position (C_18_H_34_O_2_). This fatty acid was detected in the serum EVs from healthy individuals and patients infected by pathogenic fungi (Figure 3I), with a significant increase in the concentration of EVs from patients with cryptococcosis (peak area of 1.3×10^6^), compared to healthy individuals (Figure 3J). The measured *m/z* for oleic acid was 265.2517 [M+H-H_2_O]^+^ and most of the major fragments of the compounds matched the fragmentation patterns in the GNPS database (Supplementary 3E). Putative notes detected a compound from the sphingolipids class in the EVs obtained from serum of patients infected by *C. neoformans* and *P. brasiliensis*, however further studies are required to confirm its identity and chemical structure (Supplementary 5). Thus, our results demonstrate the prevalence of lipid compounds in the serum EVs of patients with fungal infections.

### EVs from the serum and urine samples of patients with candidiasis, cryptococcosis, and paracoccidioidomycosis stimulate AMJ2-C11 cells to produce proinflammatory cytokines

To understand the inflammatory properties of EVs from patients with fungal infections, murine macrophages (AMJ2-C11) were stimulated with 10^6^ and 10^9^ EVs/mL from the serum and urine samples of patients infected by *C. albicans*, *C. neoformans*, and *P. brasiliensis* for 24 and 48 h. AMJ2-C11 cells showed a significant increase in the expression levels of interferon (IFN)-γ and tumor necrosis factor (TNF)-α at concentrations of 10^6^ and 10^9^, respectively, with serum EVs of patients with candidiasis after 24 h (Figure 4A and 4E) and 48 h (Figure 4B and 4F) compared to those in healthy individuals. Serum EVs of patients infected by *C. albicans* also exhibited significantly increased production of interleukin (IL)-6 (10^6^ and 10^9^ EVs/mL) after 24 and 48h (Figure 4I and 4J). Stimulation with serum EVs from patients with cryptococcosis showed a significant increase in the expression levels of IFN-γ (10^6^ EVs/mL) and TNF-α (10^6^ and 10^9^ EVs/mL) after 48h (Figure 4N and 4R). Moreover, analysis of cytokine production in AMJ2-C11 cells stimulated with urinary EVs revealed a significant increase in IFN-γ levels at concentrations of 10^6^ EVs/mL (24 h) and 10^9^ EVs/mL(48 h) for *C. neoformans* (Figure 4C and 4D) and TNF-α levels for patients with candidiasis compared to healthy individuals (Figure 4G and 4H). Urinary EVs of patients with cryptococcosis also promoted a significant increase in TNF-α and IL-10 levels 24 and 48 h (Figure 4K, 4L, 4O, 4P) and IL-6 levels 48 h after stimulation (Figure 4T). In turn, urinary EVs from patients infected by *P. brasiliensis* only showed an increase in TNF-α expression levels (10^9^ EVs/mL) after 24h of stimulation (Figure 4U). Our findings suggest that serum and urinary EVs of patients with candidiasis, cryptococcosis, and paracoccidioidomycosis induce murine macrophages to produce proinflammatory mediators.

**Figure 4.**
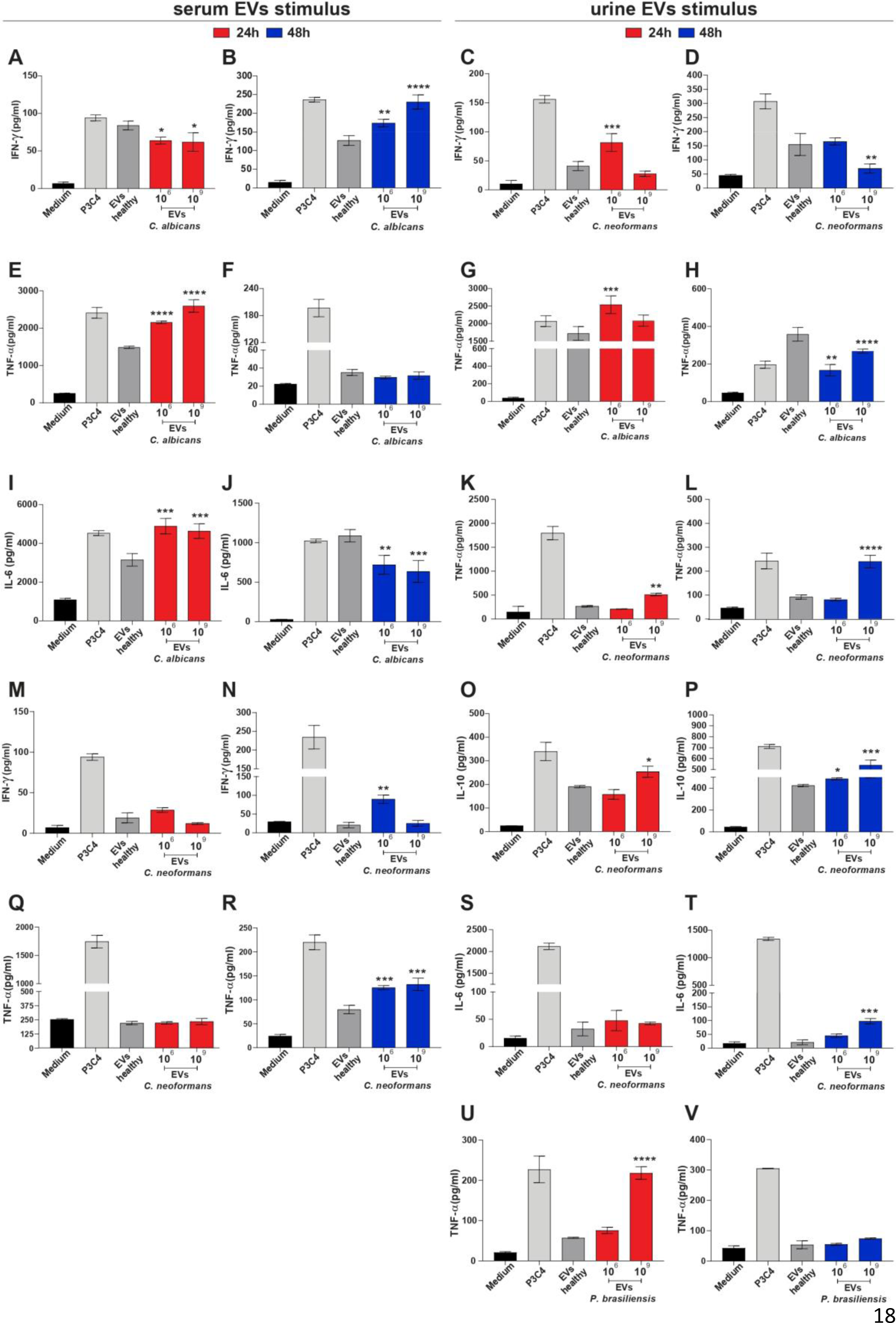
Proinflammatory mediators produced by AMJ2-C11 cells after stimulation with EVs from patients with candidiasis, cryptococcosis, and paracoccidioidomycosis. AMJ2-C11 cells (1×10^6^ cells/mL) were stimulated with Pam3CSK4 (1 μg/mL), EVs from the serum and urine samples from healthy individuals (1×10^6^ EVs/mL), and patients with fungal infections (1×10^6^ and 1×10^9^ EVs/mL), or culture medium alone. After 24 and 48 h of incubation, levels of tumor necrosis factor (TNF)-α, interferon (IFN)-γ, interleukin (IL)-6, and IL-10 in the culture supernatant were determined. Statistically significant differences in the production of the analyzed cytokines were evaluated after stimulation with serum and urine EVs from patients with candidiasis (A, B, E-J), cryptococcosis (C, D, K-T) and paracoccidioidomycosis (U and V). Values are expressed as the mean ± standard deviation (SD) compared to the levels in healthy individuals. Differences were considered significant at p < 0.05 (*); < 0.01 (**); < 0.001 (***); < 0.0001 (****).

### THP-1-derived macrophages show heterogeneous expression of cytokines after stimulation with EVs from patients infected by *C. albicans*, *C. neoformans*, and *P. brasiliensis*

THP-1 monocytic cell line can differentiate into macrophages and is often used as an in vitro model of human macrophages because of its metabolic and morphological similarities (28). Here, the same experimental conditions described above were used to verify the similarity of the immunological activity of these cells to that of murine macrophages (AMJ2-C11). THP-1-derived macrophages stimulated with EVs from the serum of patients with candidiasis exhibited increased production of IFN-γ (10^9^ EVs/mL) (Figure 5A), and IL-6 (10^6^ EVs/mL) for serum EVs of patients with cryptococcosis compared to those from the healthy individuals, after 24 h (Figure 5E). In the 48-h stimulus, EVs from patients infected by *C. neoformans* increased the production of TNF-α and IL-10 at both concentrations tested (Figure 5J and 5N). IFN-γ and TNF-α (10^6^ and 10^9^ EVs/mL) levels were significantly increased in THP-1 macrophages after stimulation with serum EVs from patients with paracoccidioidomycosis (Figure 5Q, 5R, 5U, and 5V). Stimulation with urinary EVs resulted in the heterogeneous production of cytokines in THP-derived macrophages. EVs from the urine samples of patients with candidiasis significantly increased IL-10 production (10^9^ EVs/mL) after 24 h compared to those from healthy individuals (Figure 5C). Increased production of IFN-γ (10^6^ EVs/mL) and TNF-α (10^6^ and 10^9^ EVs/mL) was observed after stimulation with EVs from the urine samples of patients with cryptococcosis in both stimulus times (Figure 5H and 5L), along with elevated IL-6 (10^9^) and IL-10 expression (10^9^) after 24 and 48h, respectively (Figure 5O, 5P, and 5T). These results show the prevalence of inflammatory mediators, mainly TNF-α and IFN-γ, in THP-1-derived macrophages. IL-10 production was also observed in the serum EVs of patients with candidiasis and serum and urinary EVs of patients with cryptococcosis.

**Figure 5.**
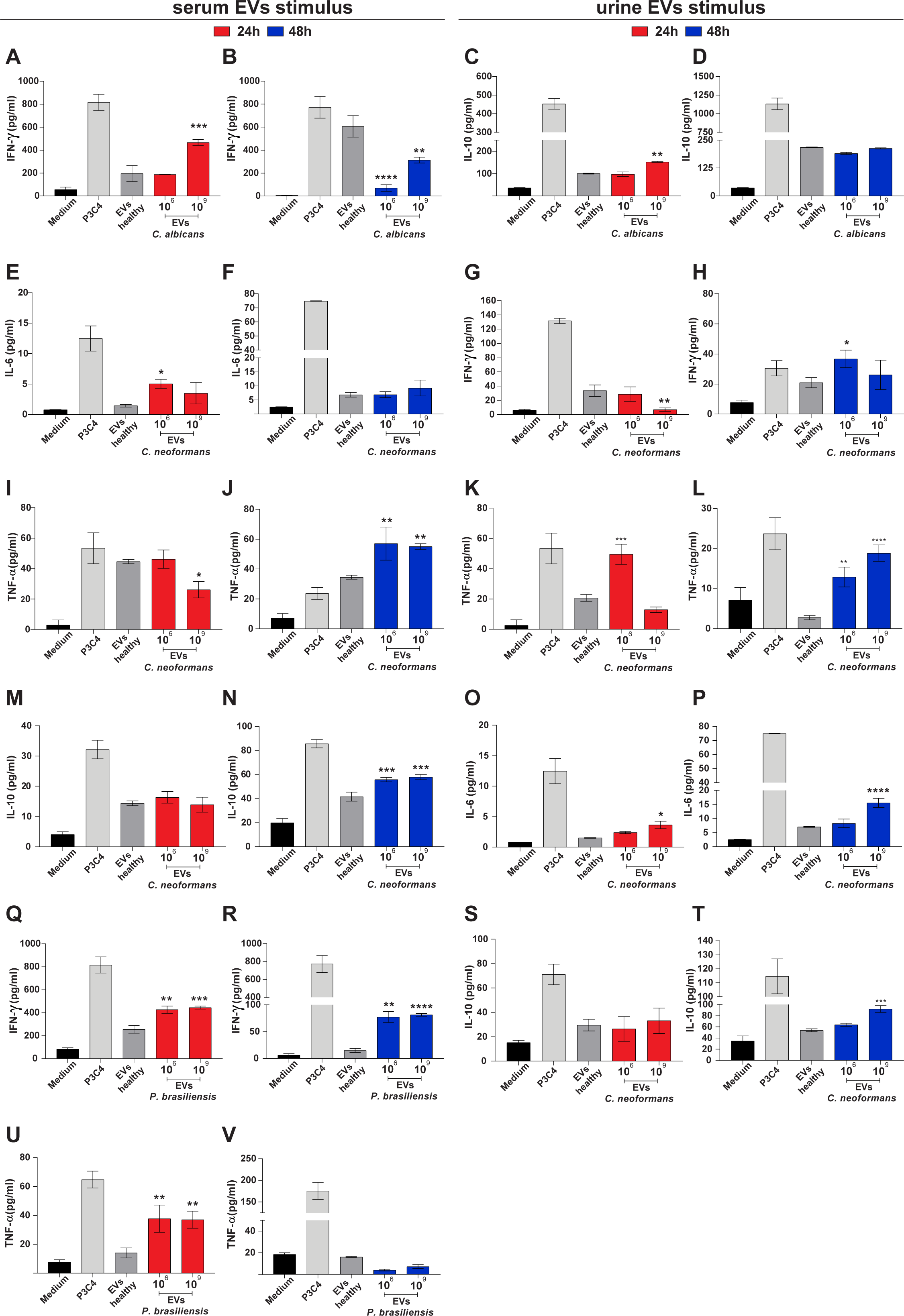
Polarization profile of macrophages derived from THP-1 human monocytes after stimulation with EVs from the serum and urine samples of patients with fungal infections. THP-1 cells (1×10^6^ cells/mL) were stimulated with Pam3CSK4 (1 μg/mL), EVs from the serum and urine samples of healthy individuals (1×10^6^ EVs/mL) and patients with fungal infections (1×10^6^ and 1×10^9^ EVs/mL), or culture medium alone. After 24 and 48 h of incubation, levels of TNF-α, IFN-γ, IL-6, and IL-10 in the culture supernatant were determined. Statistically significant differences in the production of the analyzed cytokines were evaluated after stimulation with serum and urine EVs from patients with candidiasis (A-D), cryptococcosis (E-P, S, and T), and paracoccidioidomycosis (Q, R, U, and V). Values are expressed as the mean ± SD compared to the levels in healthy individuals. Differences were considered significant at p < 0.05 (*); < 0.01 (**); < 0.001 (***); < 0.0001 (****).

### *Candida albicans* EVs induce the M1 phenotype in AMJ2-C11 macrophages

Here, the production of proinflammatory cytokines by EVs from the serum and urine samples of patients with fungal infections suggests the polarization of macrophages towards the M1 phenotype. To further analyze this process, AMJ2-C11 cells were stimulated with EVs from the serum (10^10^ EVs/mL) and urine (10^9^ EVs/mL) samples of patients for 24 h, followed by the relative quantification of transcripts of M1 (inducible NO synthase [*iNOS*]) and M2 (*Arg-1*) polarization markers. Compared to those in healthy individuals, expression levels of *iNOS* mRNA were increased by 7.7-fold in the serum EVs (Figure 6A) and 59-fold in the urinary EVs (Figure 6B) of patients with candidiasis. In terms of *Arg-1* mRNA expression, an 8-fold increase was observed in *C. albicans* (Figure 6C), whereas a 3.2-fold and 1.8-fold increase were observed in the serum and urinary EVs of *C. neoformans* (Figure 6C, and 6D). In contrast, urine samples of *P. brasiliensis* showed 1.7-fold decrease in *Arg-1* mRNA expression (Figure 6D). These results indicate that EVs of patients with fungal infections are involved in the polarization of macrophages towards the M1 phenotype.

**Figure 6.**
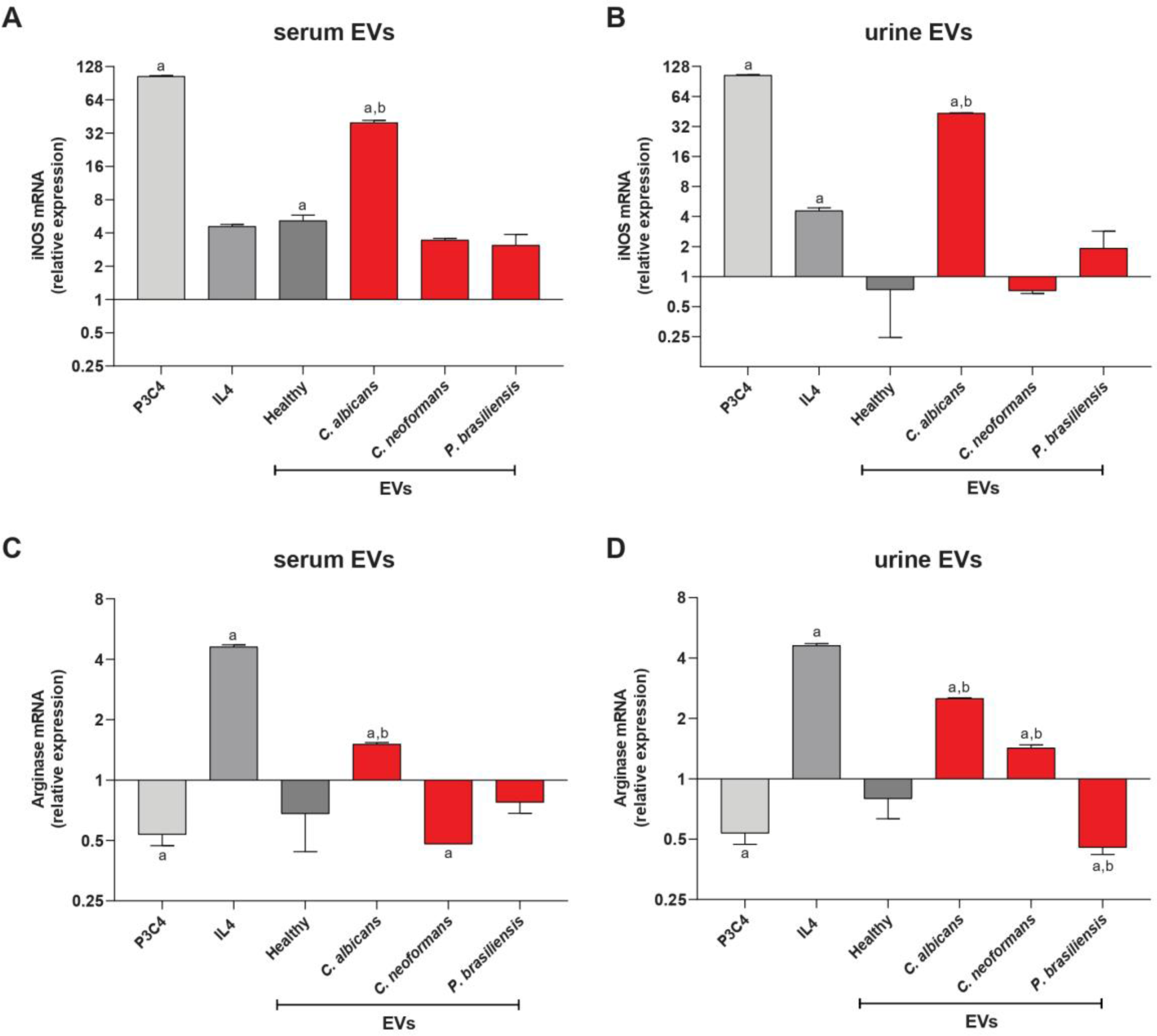
EVs of patients infected by *Candida albicans* promote the classical activation of murine macrophages. AMJ2-C11 cells (2×10^6^ cells/mL) were stimulated with EVs from the serum (1×10^10^ EVs/mL) and urine (1×10^9^ EVs/mL) samples of patients with fungal infections, EVs from the serum and urine samples of healthy individuals (1×10^6^ EVs/mL), Pam3CSK4 (1 ug/mL), IL-4 (50 ng/mL), or medium alone for 24 h at 37 °C. Relative expression levels of inducible nitric oxide synthase (*iNOS*) (A, B) and arginase 1 (*Arg-1*) (C, D) were determined via quantitative real-time polymerase chain reaction (qPCR). The results, expressed as the mean ± SD, were compared to those obtained on basal expression (statistical difference represented by the letter a) and healthy individuals (represented by the letter b).

## Discussion

In this study, we demonstrated the presence of EVs in the biological fluids, serum and urine, of patients with fungal infections caused by *C. albicans*, *C. neoformans*, and *P. brasiliensis*. The size distribution of circulating EVs in these patients was similar to that reported in previous studies in in vitro models, with a size variation of 50–100 nm, with some peaks between 350 and 500 nm for *C. albicans* (29, 30), 50–250 nm for *C. neoformans* (31), and 20–200 nm for *P. brasiliensis* (32). Collectively, these data suggest the heterogeneity of EVs among different fungal genera. The presence of EVs in the urine of patients with fungal infections (candidiasis, cryptococcosis, and paracoccidioidomycosis) also corroborates the patent by Marr and collaborators who investigated molecules present in EVs excreted in the urine of patients with invasive aspergillosis and are being explored in diagnostic methods for aspergillosis (10).

EVs from human cells are generally enriched in sphingomyelin, cholesterol, phosphatidylserine, and glycosphingolipids (33). Here, we observed the presence of cholesterol, which is important for structural rigidity and resistance to physicochemical changes, in the EVs of healthy individuals. This result was expected as cholesterol accounts for 25–50% of the lipid content in mammalian cell membranes (34). In fungal cell membranes, ergosterol is crucial for regulating membrane fluidity, protein folding, localization, and cell cycle (35). Ergosterol has been detected in the EVs of *C. albicans*, *C. neoformans*, and *P. brasiliensis* in vitro (30, 36, 37)(30, 36, 37) and not detected in healthy patients, corroborating our findings on patient samples in this study. Cholesterol/ergosterol peak area ratio may suggest the presence of EVs derived from both the host and the pathogens *C. albicans*, *C. neoformans*, and *P. brasiliensis* during fungal infection in humans.

The production of EVs by the host and fungal pathogen during infection can also be suggested due to the detection of sphingolipid class in the EVs of infected patients with candidiasis and cryptococcosis; particularly, sphingosine and phytosphingosine, compared to the EVs of healthy individuals. Sphingosine and phytosphingosine are compounds present in the cell membranes of mammals and yeast with the inclusion of specific mammalian tissues such as epidermis, small intestine, and myelin, respectively (38). Furthermore, these compounds have been identified in pathogenic strains of *C. albicans* (26) and *C. neoformans* (39), and in the EVs of *C. albicans* (40). Previous studies revealed the importance of cholesterol and sphingomyelin in controlling the host immune response in phagocytic cells (41, 42). Qureshi et al., showed the production of different species of sphingomyelin in phagocytic cells and lymphocytes and its high levels at the site of fungal infection, providing insights into the roles of sphingomyelin in host-controlled infection processes (42). However, further studies are warranted to investigate the association between sphingomyelin-derived metabolites and EVs.

The presence of oleic acid in the EVs of patients infected by *P. brasiliensis* has also been detected in vitro using the Pb18 strain (36). The significant increase in oleic acid in the EVs of patients with cryptococcosis can be explained by the fact that lipids from the host contribute to the growth, virulence, and modulation of the immune response (43). This hypothesis is supported by various in vitro and in vivo studies that showed the upregulation of genes related to the lipid metabolism of *C. neoformans* (44) and attenuated virulence of mutant strains of *C. neoformans* that do not use fatty acids as a carbon source for β-oxidation in a nutrient-scarce phagosome (45). Furthermore, Nolan and collaborators showed that the presence of oleic acid increases the replication rate of *C. neoformans*, where the fungus is able to eliminate, store, and use lipids in a beneficial way for its survival (46).

Owing to their immunobiological activity, EVs from patients with candidiasis, cryptococcosis, and paracoccidioidomycosis stimulated the murine alveolar macrophages (AMJ2-C11) and human monocyte-derived macrophages (THP-1), promoting their polarization towards a proinflammatory immune response by increasing the levels of cytokines, such as TNF-α, IFN-γ, and IL-6, and expression of the M1 gene marker characterized by *arg*. Our results are consistent with those on *C. albicans* EVs in vitro, which increased the production of NO and cytokines, such as IL-12, TNF-α, transforming growth factor (TGF)-β, IL-10, and enhanced the survival of *Galleria mellonella* upon treatment with EVs after fungal challenge (30). Here, EVs from *C. neoformans* favored the increase in the production of TNF-α and NO and phagocytosis capacity of fungal cells. However, these fungal EVs carry virulence factors, such as the capsular antigen glucuronoxylomannan, which promotes immunosuppression in macrophages with increased production of anti-inflammatory cytokines, such as TGF-β and IL-10 (16, 47). Therefore, fungal EVs derived from *C. neoformans* may play dual roles in the positive and negative stimulation of the immune response.

Here, serum and urinary EVs of patients with paracoccidioidomycosis stimulated the production of TNF-α and IFN-γ in the serum EVs of THP-1 macrophages. The classical inflammatory activation profile (M1) promoted by *P. brasiliensis* EVs is characterized by the production of NO and proinflammatory cytokines (TNF-α, IL-6, and IL-12p70). Incubation of alternatively activated macrophages (M2), which were subsequently treated with EVs from this fungus, changed their phenotype to an M1 profile (15). Furthermore, EVs from the attenuated variant of the Pb18 strain significantly increased the expression of inflammatory mediators (NO, TNF-α, and IL-6) compared to those from the virulent strain (48). To the best of our knowledge, this is a pioneering study on the identification and characterization of EVs from the clinical samples of patients with fungal infections caused by *C. albicans*, *C. neoformans*, and *P. brasiliensis*, providing new insights into the roles of EVs in humans and their possible use in therapeutic strategies for systemic mycoses.

## Materials and Methods

### Clinical samples

Blood and urine samples from healthy individuals and patients with fungal infections caused by *Candida albicans*, *Cryptococcus neoformans*, and *Paracoccidioides brasiliensis* were obtained from Hospital das Clinicas, Ribeirao Preto Medical School, University of Sao Paulo, SP, Brazil (HC-FMRP, USP). Sample collection was carried out in collaboration with Prof. Dr. Roberto Martinez and Prof. Dr. Silvana Maria Quintana (HC-FMRP). Blood samples were collected in a collection tube with gel and clot activator (Firstlab) and centrifuged at 2000×g for 10 min (Thermo Fisher Sorvall Legend RT Centrifuge) to obtain serum. Urine samples were collected from the midstream of the first-morning urine in a sterile universal collector (Fisrtlab). The procedures performed with patients follow the principles proposed by the National Research Ethics Commission and the Research Ethics Committee (protocol HCRP 4.096/2012).

### Inclusion and exclusion criteria of the patients

Female and male patients, without a defined age range, were selected for the clinical study. The inclusion criterion was the presence of fungal infections caused by *C. albicans*, *C. neoformans*, and *P. brasiliensis*, in the acute phase. The clinical samples of candidiasis patients consisted of vulvovaginal manifestation. Exclusion criteria were the presence of human immunodeficiency virus (HIV), and other bacterial acute infections, as well as previous treatment for fungal infections.

### Murine alveolar macrophage cell line (AMJ2-C11)

The murine alveolar macrophage cell line (AMJ2-C11) was obtained from the Rio de Janeiro Cell Bank (BCRJ). The cells were maintained in Dulbecco’s modified Eagle’s medium (DMEM) with high glucose (Gibco-Thermo Fisher Scientific) supplemented with fetal bovine serum (10%), sodium bicarbonate, and penicillin/streptomycin (1%), and incubated in a CO_2_ (5%) at 37°C. Cells were cultivated until confluence and detached from the flask bottom with PBS pH 7.4 (Gibco, Thermo Fisher) and 0.5M EDTA pH 8.0 (Invitrogen, Thermo Fisher Scientific). Cells were plated in 24 or 48-well culture plates, depending on the assay performed.

### Human THP-1-derived macrophages

The human monocytic cell line THP-1 (American Type Culture Collection ATCC TIB202) is derived from an acute monocytic leukemia cell line. The cells were maintained in a suspension culture dish (Corning Inc. Sigma Aldrich) with RPMI medium 1640 medium (Gibco-Thermo Fisher Scientific) supplemented with fetal bovine serum (10%), sodium bicarbonate, and penicillin/streptomycin (1%), and incubated in a CO_2_ (5%) at 37°C. THP-1 monocytes were differentiated into macrophages as previously described (28). THP-1 cells were stably differentiated into macrophages by phorbol 12-myristate 13-acetate (PMA) (Sigma-Aldrich) at 5 ng/ml over 48h. Cells differentiated into macrophages were plated in 6-well plates for the assay.

### Standardization of extracellular vesicle (EVs) extraction in clinical samples

EVs were isolated as previously described (37, 49), with some modifications. First, the collected blood samples were centrifuged at 2,000×g for 10 minutes (Thermo Fisher Sorvall Legend RT Centrifuge) to obtain the serum. Serum (2 ml) and urine (15 ml) samples were subjected to centrifugation at 2,000×g, 15 min, at 4°C to remove cells and debris (Thermo Fisher Sorvall Legend RT Centrifuge). In turn, serum samples were centrifuged at 10,000×g, 30 min, 4°C (Thermo Fisher Sorvall Legend RT Centrifuge), while urine samples were concentrated through a polystyrene membrane in an Amicon ultracentrifugation system with a 100kDa cutoff (Millipore, Billerica, MA, USA) and then centrifuged at 15,000×g, 15 min, 4°C (Thermo Fisher Sorvall Legend RT Centrifuge). Subsequently, the supernatants obtained from serum and urine samples were filtered through a 0.22 μm filter and ultracentrifuged at 100,000×g, 1 hour, 4°C (Optima TLX Ultracentrifuge, Beckman Coulter). The pellet was resuspended in 200 µL of ultrapure water.

### Nanoparticle tracking analysis (NTA)

Size distribution analysis and quantification of EV preparations were performed on a NanoSight NS300 (Malvern Instruments, Malvern, UK) equipped with rapid video capture and particle tracking software. Vesicles purified from clinical samples were diluted in 1 mL of ultrapure water. Each sample was injected into the sample cubicle of the device for analysis. Fluorescence scattering and capture settings (such as focus, camera, and gain settings) were optimized to make particle tracks visible, and then measurements were obtained in triplicate and analyzed using NanoSight NS300 software (version 3.2.16). Data on EV sizes were expressed as the calculated means ± SD of size distribution.

### Transmission electron microscopy (TEM)

Serum and urine EV samples from healthy individuals and patients with candidiasis, cryptococcosis, and paracoccidioidomycosis were incubated with glutaraldehyde 2% plus paraformaldehyde 2% in 0.1M sodium cacodylate buffer (pH 7.4) for 4 hours at a temperature of 2°C to 4°C and subsequently ultracentrifuged at 100,000×g at 4°C for 1 hour. The pellet obtained from the samples was resuspended in 200 μl of sodium cacodylate buffer. Images were captured with a JEOL JEM 100CXII transmission electron microscope with a Hamamatsu ORCA-HR digital camera.

### Western blotting of classical EVs markers

Western blotting of EVs from the serum and urine of healthy individuals and patients with candidiasis, cryptococcosis, and paracoccidioidomycosis was performed as previously described (50), with some modifications. Serum and urine EVs were lysed with RIPA buffer containing protease inhibitors. The samples were sonicated for 1 min at 60% amplitude (10 on and 10 off pulses), centrifuged at 15,000×g, for 20 min at 4°C, and the supernatants were collected for quantification. Protein concentration was determined by the Pierce BCA Protein Assay Kit method (Thermo-scientific, catalog numbers 23225 and 23227), separated by SDS-PAGE, and transferred to nitrocellulose membranes for 1 hour. The membranes were blocked with 5% BSA in 0.05% Tween/TBS and incubated with the specific antibodies. ALIX (E47TU-Cell Signaling), Anti-CD63 (SAB4301607-Sigma Aldrich), and Anti-CD9 (EPR23105121-Abcam) were used at a dilution of 1:1000. The protein-antibody complex was detected using ECL Western blot detection reagents (GE Lifesciences), and signals were detected using a CCD camera (Image Quant LAS 4000 mini, Uppsala, Sweden).

### Mass spectrometry (MS)

Extracellular vesicles obtained from the serum and urine of healthy individuals and patients diagnosed with candidiasis, cryptococcosis, and paracoccidioidomycosis were subjected to extraction using 200 μL of methanol, followed by drying in a stream of N2. The extracts obtained were prepared at a concentration of 1 mg/mL and subjected to analysis via liquid chromatography-high resolution mass spectrometry/mass spectrometry (LC-HRMS/MS) using a (Thermo Scientific QExactive® Quadrupole-Orbitrap Hybrid instrument). The analyzes were conduced in both positive and negative modes with a *m/z* range of 100-1500, a capillary voltage of 3.4 kV, an inlet capillary temperature of 280 °C, a voltage of 100 V. Chromatographic separation employed a Thermo Scientific Accucore C18 2.6 μm (2.1 mm x 100 mm) column. The stationary phase consisted of MilliQ water with 0.1% formic acid (A) and acetonitrile with 0.1% formic acid (B). Chromatographic separation was performed using the following gradients: 0-5 min, 5% B; 5-10 min, 40% B; 10-12 min, 45% B; 12-18 98% B; 18-20 min, 98% B; 20-22 min, 5% B and 22-24 min 5% B. The injection volume of each sample was standardized at 10 μL. MS data were collected in data-dependent acquisition (DDA) mode, capturing MS/MS spectra of the five most intense ions in 5-s windows. The Orbitrap analyzer resolution was set as 70,000, with an Automatic Gain Control (AGC) target of 1e6 and a maximum injection time (IT) of 100ms. MS/MS spectra were obtained in centroid mode, maintaining a resolution of 35,000, with a target AGC of 1e5 and a maximum IT of 100 ms. Collision energies in normalized collision energy (NCE) mode were applied at 20, 35, and 45 eV. MS data were processed with Xcalibur software (version 3.0.63) developed by Thermo Fisher Scientific.

### Molecular Networks and Metabolomics Analysis

The raw files (.raw) were converted via MSConvert (51) to the mz.XML format and analyzed in the Mzmine 3.9.0 software, the processing parameters used are in the table 1. The .mgf files and the quantification tables obtained by Mzmine were used to perform Feature Based Molecular Networking (FBMN), where the fragmentation tolerance parameters of the library data were 0.02 Da, and the cosine score for the chromatograms that presented corresponding peaks was 0.7 (70%). Pre-established libraries were used for putative annotations according to the feature table generated by MZMine 3.9.0 (52). The annotated compounds were classified according to (53): Compounds confirmed by comparison of MS/MS profiles with reference standards (In this study reference standards for Cholesterol, Ergosterol and D-erytro sphinganine were used) – Level I Annotation; Compounds annotated through comparison of the MS/MS fragmentation profile with spectra present in the GNPS libraries – Annotation Level II; Compounds annotated through simulated spectra used were generated by SIRIUS 5.7.0-Annotation Level III; Compounds not annotated, but with molecular formula unequivocally determined through information from LC-HRMS analyzes – Level IV Annotation. The molecular networks, as well as the annotations made against the GNPS databases, are available at: https://gnps.ucsd.edu/ProteoSAFe/status.jsp?task=3396730133064ed6a81bb5cfa59a0208. The FBMN molecular networks were visualized using Cytoscape (v.3.10.1), in total 533 compounds were annotated according to the GNPS MS/MS libraries. All MS1 and MS/MS spectra data, together with their metadata in mzXML format, have been deposited in an open format in the public data repository GNPS/MassIVE (54). Finally, the areas of the differential ions noted through univariate and multivariate comparisons were calculated manually using Thermo Xcalibur 3.0.63 software (Copyright 1988-2013 Thermo Fisher Scientific Inc.), and correlation plots of the calculated areas were constructed to analyze the abundances of these compounds in the different sample groups, using Prisma 8.0.1 software (Graph Pad Software, San Diego, CA).

**Table 1.** MS1 and MS2 preprocessing parameters.

| Pre-processing Chromatogram | Parameter Value |
| --- | --- |
| Crop retention time | 0.3-24.00 min |
| RT Tolerance (intra-sample) | 0.10 min |
| RT Tolerance (sample-to-sample) | 0.4 min |
| Pre-processing MS | Parameter Value |
| MS noise level | 1.0 x 10 |
| MS noise level | 3.0 x 10 |
| Minimum feature height | 5.0 x 10 |
| *m/z* tolerance (scan-to-scan) | 5 ppm |
| *m/z* tolerance (intra-sample) | 3 ppm |

### Immunobiological activity of murine and human macrophages after incubation with EVs from clinical samples

Murine alveolar macrophages (AMJ2-C11) and human macrophages (THP1) were plated in 48- and 6-well plates, respectively (1.5×10^6^ cells/ml), and maintained at 37°C, 5% CO_2_, overnight. AMJ2-C11 and THP1 cells were stimulated with pools of EVs from the serum and urine of patients with candidiasis, cryptococcosis, and paracoccidioidomycosis at concentrations of 1×10^6^ and 1×10^9^ EVs/ml. As a control, EVs from serum and urine from healthy individuals were used at a concentration of 1×10^6^ EVs/ml, Pam3CSK4 (1μg/ml), and culture medium alone. The cells were incubated for 24 and 48 hours at 37°C with 5% CO_2_. The supernatant was collected to measure TNF-α, IFN-γ, IL-6, and IL-10 using an enzyme-linked immunosorbent assay (ELISA) with a BD Biosciences kit (San Diego, CA, USA), as described by the manufacturer. The absorbance value was determined at 450 nm on a spectrophotometer for microplates (SpectraMax 190; Molecular Devices).

### Quantitative Reverse Transcription PCR (qPCR)

AMJ2-C11 were plated in 24-well plates (2×10^6^ cells/ml) and stimulated with EVs from serum (1×10^10^ EVs/ml) and urine (1×10^9^ EVs/ml) from patients with fungal infections, EVs from serum and urine from healthy individuals (1×10^6^ EVs/ml), Pam3CSK4 (1ug/ml), IL-4 (50ng/ml), or medium alone for 24 h at 37°C in a 5 % CO_2_ atmosphere. The supernatant was discarded, and the cells were used for RNA extraction using the Illustra RNAspin Mini RNA isolation kit (GE Healthcare) following the manufacturer’s instructions. RNA concentration and quality were measured using a nanophotometer (Implem). The reaction for cDNA synthesis was performed using 1 μg of RNA, following the instructions available in the High-Capacity cDNA Reverse Transcription Kit (Applied Biosystems®). Real-time Polymerase Chain Reaction (qPCR) was conducted in a StepOnePlus™ Real-time PCR system (Applied Biosystems®) to analyze the relative expression of the genes of inducible nitric oxide synthase (*iNOS*) and arginase-1 (*Arg-1*). The β-actin gene (*Actb*) was used as an endogenous reference control. The primer sequences for these macrophage-polarization markers are listed in Table 2. The conditions used in the qPCR reactions followed the protocol available in the qPCRBIO SyGreen Mix kit (PCR Biosystems®). All samples were analyzed in triplicate. Relative expressions were calculated by comparing Threshold Cycle (Ct) values of β-actin and targeted genes according to the 2^-ΔΔCT^ method described by (55).

**Table 2.**
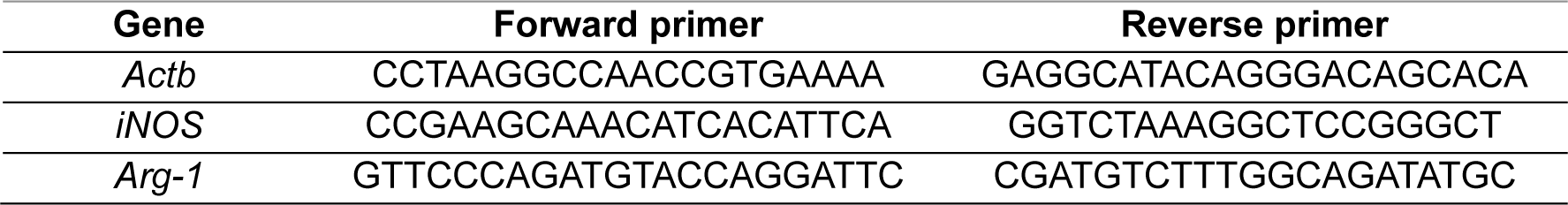
Nucleotide sequences of primer sets used to amplify the genes using qPCR.

### Statistical analysis

The results were expressed as the mean ± SD (standard deviation). Statistical analyses of data with more than three experimental groups were performed using the One-Way ANOVA test, Sidak’s multiple comparisons test with a single pooled variance, using the Prisma 8.0.2 program (Graph Pad Software, San Diego, CA). Statistical differences are p < 0.05.

## Acknowledgments

We thank Carlos Alberto Vieira, Maria Dolores Seabra Ferreira, and José Augusto Maulin for technical support. This research was funded by Fundação de Amparo à Pesquisa do Estado de Sao Paulo [2020/03215-1; 2021/06794-5], CNPq [Conselho Nacional de Desenvolvimento Científico e Tecnológico], and CAPES [Coordenação de Aperfeiçoamento de Nível Superior]. Institutional Review Board Statement: The study was conducted according to the principles proposed by the National Research Ethics Commission and approved by the Research Ethics Committee of the Ribeirao Preto Medical School at the University of Sao Paulo, Protocol HCRP 4.096/2012.

## Author Contributions

Conceptualization: C.P.R., M.L.R., R.M., and F.A.; methodology: C.P.R., P.W.S., R.A.P, V.C.S, J.C.J.B., and T.P.F; validation: T.P.F. and F.A.; formal analysis: C.P.R., R.A.P., J.C.J.B., and T.P.F.; image formatting: C.P.R. and P.W.S.; clinical samples: C.P.R., F.C.P.V., E.T., R.C., G.P., S.M.Q. and R.M.; resources: F.A.; data curation: C.P.R., R.A.P., J.C.J.B,, T.P.F., and F.A.; writing— original draft preparation, C.P.R. and F.A.; writing—review and editing, C.P.R. and F.A.; visualization: C.P.R.; supervision: F.A.; project administration: F.A. All authors have read and agreed to the published version of the manuscript.

